# FGF21, a liver hormone that inhibits alcohol intake in mice, increases in human circulation after acute alcohol ingestion and sustained binge drinking at Oktoberfest

**DOI:** 10.1101/264234

**Authors:** Susanna Søberg, Emilie S. Andersen, Niels B. Dalgaard, Ida Jarlhelt, Nina L. Hansen, Nina Hoffmann, Tina Vilsbøll, Anne Chenchar, Michal Jensen, Trisha J. Grevengoed, Sam A.J. Trammell, Filip K. Knop, Matthew P. Gillum

## Abstract

**Objective:** Excessive alcohol consumption is a leading cause of global morbidity and mortality. However, knowledge of the biological factors that influence ad *libitum* alcohol intake may be incomplete. Two large studies recently linked variants in the *KLB* locus with levels of alcohol intake in humans. *KLB* encodes ß-klotho, co-receptor for the liver-derived hormone fibroblast growth factor 21 (FGF21). In mice, FGF21 reduces alcohol intake, and human *Fgf21* variants are enriched among heavy drinkers. Thus, the liver may limit alcohol consumption by secreting FGF21. However, whether full-length, active plasma FGF21 (FGF21 (1-181)) levels in humans increase acutely or sub-chronically in response to alcohol ingestion is uncertain.

**Methods:** We recruited 10 healthy, fasted male subjects to receive an oral water or alcohol bolus with concurrent blood sampling for FGF21 (1-181) measurement in plasma. In addition, we measured circulating FGF21 (1-181) levels, liver stiffness, triglyceride, and other metabolic parameters in three healthy Danish men before and after consuming an average of 22.6 beers/person/day (4.4 g/kg/day of ethanol) for three days during Oktoberfest 2017 in Munich, Germany. We further correlated fasting FGF21 (1-181) levels in 49 healthy, non-alcoholic subjects of mixed sex with self-reports of alcohol-related behaviors, emotional responses, and problems. Finally, we characterized the effect of recombinant human FGF21 injection on ad *libitum* alcohol intake in mice.

**Results:** We show that alcohol ingestion (25.3 grams or ~2.5 standard drinks) acutely increases plasma levels of FGF21 (1-181) 3.4-fold in fasting humans. We also find that binge drinking for three days at Oktoberfest is associated with a 2.1-fold increase in baseline FGF21 (1-181) levels, in contrast to minor deteriorations in metabolic and hepatic biomarkers. However, basal FGF21 (1-181) levels were not correlated with differences in alcohol-related behaviors, emotional responses, or problems in our non-alcoholic subjects. Finally, we show that once-daily injection of recombinant human FGF21 reduces ad *libitum* alcohol intake by 21% in mice.

**Conclusions:** FGF21 (1-181) is markedly increased in circulation by both acute and sub-chronic alcohol intake in humans, and reduces alcohol intake in mice. These observations are consistent with a role for FGF21 as an endocrine inhibitor of alcohol appetite in humans.

## 1. INTRODUCTION

Alcohol abuse is implicated in more than 2.3 million deaths worldwide every year, and is the ninth largest contributor to the global burden of disease in terms of disability-adjusted life years (DALYs) [1]. Moreover, longitudinal studies demonstrate that alcohol abuse precedes poor outcomes, suggesting that it is the cause, rather than a consequence, of these problems [2]. However, drugs that adequately control alcohol use disorders have yet to be developed, in part because the biology that motivates alcohol consumption is incompletely understood [3].

Recently, a GWAS including >100,000 subjects of European descent associated rs11940694 in the *KLB* locus with levels of alcohol intake in non-alcoholic subjects [p=9.2 × 10^−12^] [4]. This result was replicated in an additional 112,117 subjects from the UK Biobank that also linked rs11940694 [p=8.4 × 10^−19^] as well as rs9991733 [P=3.80 × 10^−9^] in *KLB* with alcohol drinking in humans [5]. The*KLB* gene encodes β-klotho, an essential co-receptor *in vivo* for the metabolic hormones FGF19 and FGF21 [6].

Circulating FGF21 is mainly derived from the liver and is best known for its beneficial effects on glucose, lipids, and body weight when given pharmacologically to rodents [7, 8]. In humans, FGF21 has limited effects on glycemia, but does improve lipid homeostasis and weight, suggesting that some, but not all, of its rodent functions may be conserved [9, 10]. In mice, alcohol increases circulating FGF21 levels [11, 12] and FGF21 overexpression or continuous infusion by minipump reduces alcohol and sweet intake via a ß- klotho-dependent cascade in the central nervous system [4, 13]. In addition, a bispecific antibody that activates the β-klotho/FGFR1 complex reduces alcohol and saccharin intake in mice [14]. In favor of a conserved role for FGF21 as a negative regulator of alcohol and sugar intake in humans, the *Fgf21* rs838133 A-allele is enriched among the top tertile of alcohol drinkers and sweet consumers in the Danish Inter99 cohort [15]. Moreover, it was recently reported that oral ethanol markedly increases circulating total FGF21 levels in non-fasted human subjects [16]. Together, these observations suggest that alcohol ingestion increases hepatic secretion of FGF21, which then acts via the β-klotho/FGFR1 complex to suppress further alcohol intake.

To test this prediction, we assessed plasma FGF21 (1-181) levels in fasted humans after acute and sub-chronic alcohol ingestion, and measured the effects of bolus human FGF21 administration on alcohol preference in mice. We further correlated fasting FGF21 (1-181) levels with self-reported alcohol-related behaviors, emotional responses, and problems in a cohort of 49 healthy human subjects.

## 2. MATERIALS AND METHODS

### 2.1.1 Subjects for the acute and questionnaire alcohol studies

Male subjects age 18-30 years, with a body mass index (BMI) of 19-25 kg/m^2^ and without any medical conditions were recruited for participation by advertisement. Individuals with chronic diseases, history of alcohol abuse, and current or former smoking were excluded. Additionally, individuals who were on a diet, were taking dietary supplements, had a history of eating disorders, recently experienced weight loss (> 3 kg within 3 months), or used daily medications were excluded from participation. The study was approved by the Scientific Ethics Committee of the Capital Region of Denmark (H-16038581). All subjects provided written and informed consent prior to participation in the study, which was performed in accordance with the principles enumerated in the Declaration of Helsinki II.

The 10 subjects selected for the acute alcohol (ethanol) study underwent a pre-examination that included baseline laboratory tests (after a 12 hour overnight fast), medical history, and physical examination. A 2-hour oral glucose tolerance test (OGTT) was performed to confirm that none of the subjects had impaired glucose tolerance (9-11 mmol/l after two hours) or were insulin resistant (plasma insulin levels above normal: 10-125 pmol/l). Subjects were instructed to refrain from alcohol and caffeine intake 48 hours prior to participation in the study and none of the subjects had any significant alcohol drinking at least one week prior to the study day.

Recruitment of subjects for the alcohol questionnaire cohort (whose baseline FGF21 levels were correlated with responses to the Alcohol Use Disorder Test (AUDIT) and Alcohol Use Disorders Identification Test-E (ALCOHOL-E), combined with those of subjects from the ethanol study) has been described elsewhere [15]. In total, 39 subjects of the 52 from this cohort, which consisted of young, healthy subjects of mixed sex, answered the alcohol questionnaires that were sent by e-mail. Subjects were advised to sit alone while answering the questionnaires, not be under the influence of alcohol, and to answer as honestly as possible with the understanding that their responses were anonymous.

### 2.1.2 Oktoberfest cases

Three healthy Danish men (aged 42 years, BMIs of 23.8, 21.4, 28.3 kg/m^2^) with normal everyday alcohol habits contacted our clinic for a health examination before three days of Oktoberfest participation in Munich, Germany. Upon their return they asked for a reevaluation of their metabolic health (32 hours after consuming the last unit of alcohol at Oktoberfest). All agreed to have their cases reported anonymously.

### 2.2.1 Acute alcohol study

On the study day, subjects in the acute ethanol cohort met after an overnight fast (12 hour) and 48 hour abstention from physical activity and caffeine intake. Furthermore, subjects were instructed not to consume alcohol one week prior to the study day. Blood samples were collected in the fasting state from an antecubital vein at −10 and 0 minutes and 15, 30, 60, 90, 120, 180 and 240 minutes after ingestion of 280 ml of either tap water or ethanol mixed with tap water (8 centiliters of 40% ethanol in 200 ml tap water). Subjects were randomized to either the water control group or the alcohol group using a computer randomization system at www.easytrial.net. During the study day the subjects were resting in a hospital bed and instructed to relax but not sleep.

### 2.2.2 Oktoberfest case examinations

Subjects were examined in the morning after an overnight (10 hour) fast. Body weight, height, body composition (by bioimpedance) and hepatic stiffness (by fibroscanning) were evaluated and blood and urine was sampled. Insulin resistance was measured by the homeostatic model assessment (HOMA-IR) and homeostatic model assessment 2 (HOMA-2IR) with the HOMA2-IR being a computerized model recalibrated to the modern insulin assays. HOMA-IR is calculated using the fasting plasma glucose and the fasting insulin levels divided by 22.5 (HOMA-IR= [FPG*FPI]/22.5).

### 2.3. Alcohol questionnaires

Subjects in the acute ethanol cohort (n=10) and the sweet-liking cohort (n=39) answered two questionnaires concerning their alcohol habits. One of the questionnaires, AUDIT, is a validated questionnaire, used to identify patients at risk of alcohol-related health problems or alcohol dependence. The questionnaire consists of 10 questions divided into three categories; *Alcohol intake, Alcohol dependence,* and *Harmful alcohol use.* The score for all 10 questions is aggregated and the total AUDIT score can range from 0-40. The score is interpreted as follows: a score of 7 indicates an alcohol problem; a score of 8-15 indicates a large amount of consumption that can be remedied by short intervention; a score of 16-19 suggests harmful consumption, which requires short intervention and/or medical treatment; a score of ≥ 20 strongly suggests addiction, but addiction cannot be excluded if the score is below this limit.

The second alcohol-questionnaire, "Alcohol-E" (Alcohol Use Disorders Identification Test-E), was originally developed in Sweden to identify alcohol use disorders. In this study, we used the Danish version of Alcohol-E. Alcohol-E provides a structured view of a person’s alcohol use. The questionnaire consists of 4 sections with the following questions: *1. This often I drink these alcoholic drinks?* (e.g. beer, strong beer, wine and alcoholic soda-drinks); *2. What is positive for you when using alcohol?* (e.g. sleep better, get happy, feel strong and getting better at contacting other people); *3. What is negative for you when using alcohol?* (e.g. become anxious, have suicidal thoughts, have less contact with friends, experience low libido); *4. What do you think about alcohol?* (e.g. do you enjoy drinking alcohol, do you feel it is important to change your alcohol use, and have you the past year been concerned about your alcohol use?). In this study, we used the “Alcohol-E” questionnaire to get a deeper insight into which alcoholic drinks our subjects preferred and to achieve more knowledge on the negative and positive effects of their drinking habits. The results from the questionnaire were calculated by dividing the mean score for each section by the number of questions in that section.

### 2.4. FGF21 measurements

Blood samples used for FGF21 (1-181) measurements were put on ice immediately after collection and separated by centrifugation for 15 minutes at 3000 rpm. Plasma samples were subsequently stored in aliquots at −20 to −80 °C pending further analysis. FGF21 levels were measured using a commercially available enzyme-linked immunosorbent assay (ELISA) (Eagle Biosciences, F2131-K01) that detects full-length, active FGF21 (1-181), but not FGF21 fragments generated via recently described cleavage events at the N-terminus by dipeptidyl peptidase 4 (DPP-4), or at the C-terminus by fibroblast activation protein (FAP) that could lead to overestimates of the amount of FGF21 in circulation [17, 18]. The standard curve range for the assay is 32.5-2000 pg/ml, with a LoD of 1.7 pg/ml. FGF21 was assayed in duplicate and calculated using the mean of replicates. Total FGF21 was also measured in duplicate in the Oktoberfest group using a kit that detects total FGF21 (R&D Systems, DF2100). The LoD for this assay is 4.67 pg/ml and a standard curve ranging from 31.3-2000 pg/ml.

### 2.5. General hormone analyses

All subjects in the acute alcohol study attended a pre-examination day with routine blood samples to screen for diseases before they were included in the experiment. Measurements of hemoglobin and glycosylated hemoglobin (Hb1Ac) and plasma levels of insulin, C-peptide, glucose, cholesterol, triglycerides, ALAT, ASAT, thyroid hormones, thyrotropin, alkaline phosphatase, carbamide, and urea were performed using standard techniques at the Department of Clinical Biochemistry, Rigshospitalet, University of Copenhagen, Copenhagen, Denmark or at the Department of Clinical Biochemistry, Gentofte Hospital, University of Copenhagen, Hellerup, Denmark. Blood samples for insulin, C-peptide, and glucose were immediately spun at 4°C at 3000 rpm for 15 min, and were stored at 2-4°C until analysis the same day. Plasma insulin was analyzed by electrochemiluminescent immunoassay (Cobas, Roche) and C-peptide by sandwich electrochemiluminesence-immunoassay (ECLIA).

### 2.6. Mouse alcohol preference test

Eighteen male C57BL/6N mice (8-10 weeks old; Taconic, Denmark) were singly housed and given the choice between water and a 4% ethanol solution. Every day, the consumption of both was determined by weighing the drinking bottles. A stable, 4-day, pre-treatment baseline intake was established for each mouse. Every other day, the position of the two drinking bottles was reversed to ensure that there was no side-preference bias for any of the mice. Half of the mice were treated with FGF21, through intraperitoneal injection with a solution of FGF21 (10 mg/ml in PBS, a 1 mg/kg dose) for 3 days. The other half received injections of PBS during the same period.

### 2.7. Statistical analysis

All analyses were performed using the Statistical Package for SAS 9.1 (SAS Institute, Cary NC, USA) or GraphPad Prism v6.0 (GraphPad Software, La Jolla, CA). Data are presented as mean and standard deviations for normally distributed variables. Relevant statistical test, sample size and significance level for each analysis are stated in figure legends. p-values below 0.05 where Benjamani-Hochberg Q<0.05—if relevant due to multiple comparison—were considered statistically significant.

## 3. RESULTS

### 3.1 Alcohol ingestion acutely increased levels of active FGF21 (1-181) in human plasma

Ten healthy, fasted male subjects were randomly assigned to groups similar in anthropometric and metabolic variables at baseline (Table 1). After an overnight fast and basal blood sampling, all participants received a single oral bolus of either water or alcohol, and blood samples were collected for the next four hours while participants did not eat or drink, since nutrients influence hepatic FGF21 production in a complex manner [19, 20].

After drinking water or 8 centiliters 40% alcohol (25.3 grams pure ethanol) diluted in water to an equal total volume, blood alcohol level (BAC) increased only in the alcohol group as anticipated, rising to a peak of 0.043% at 30 minutes, and returning to zero after three hours (Supplementary Figure 1). Because increases in blood glucose can increase circulating FGF21 levels [21], we also measured plasma glucose, insulin and C-peptide during the experiment. Although differences in insulin or C-peptide were not observed between groups, their absolute levels decreased over time, consistent with the increased period of fasting. However, plasma glucose levels were significantly lower in the alcohol group after 90 minutes, and remained lower for the duration of the experiment (Supplementary Figure 1).

**Table 1.**
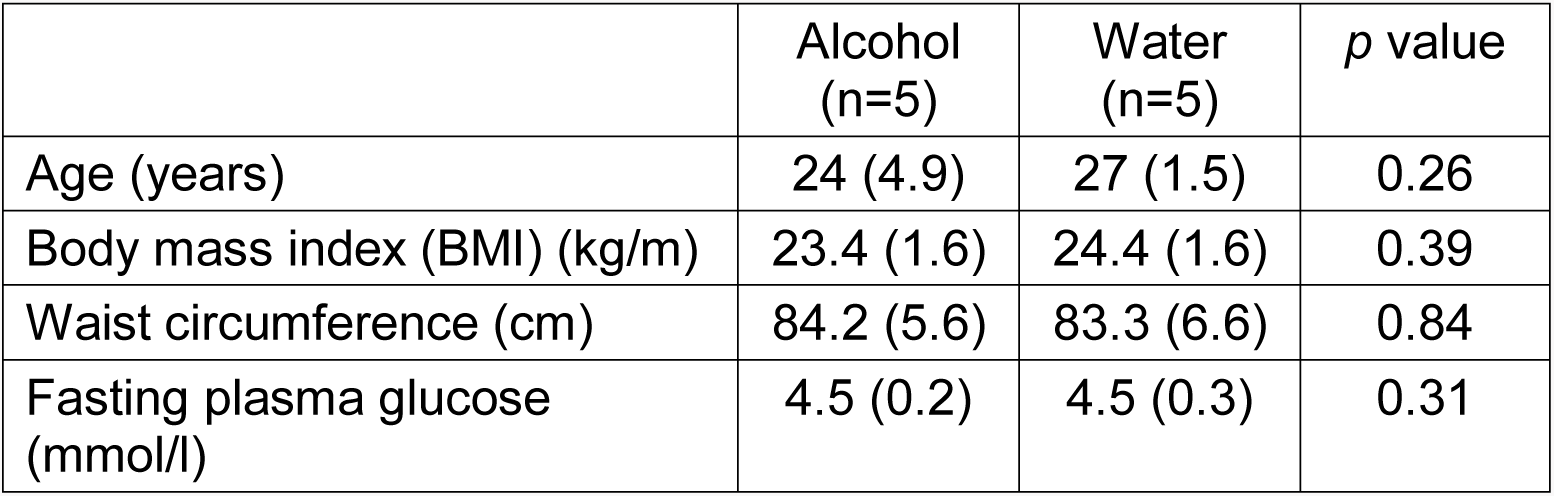
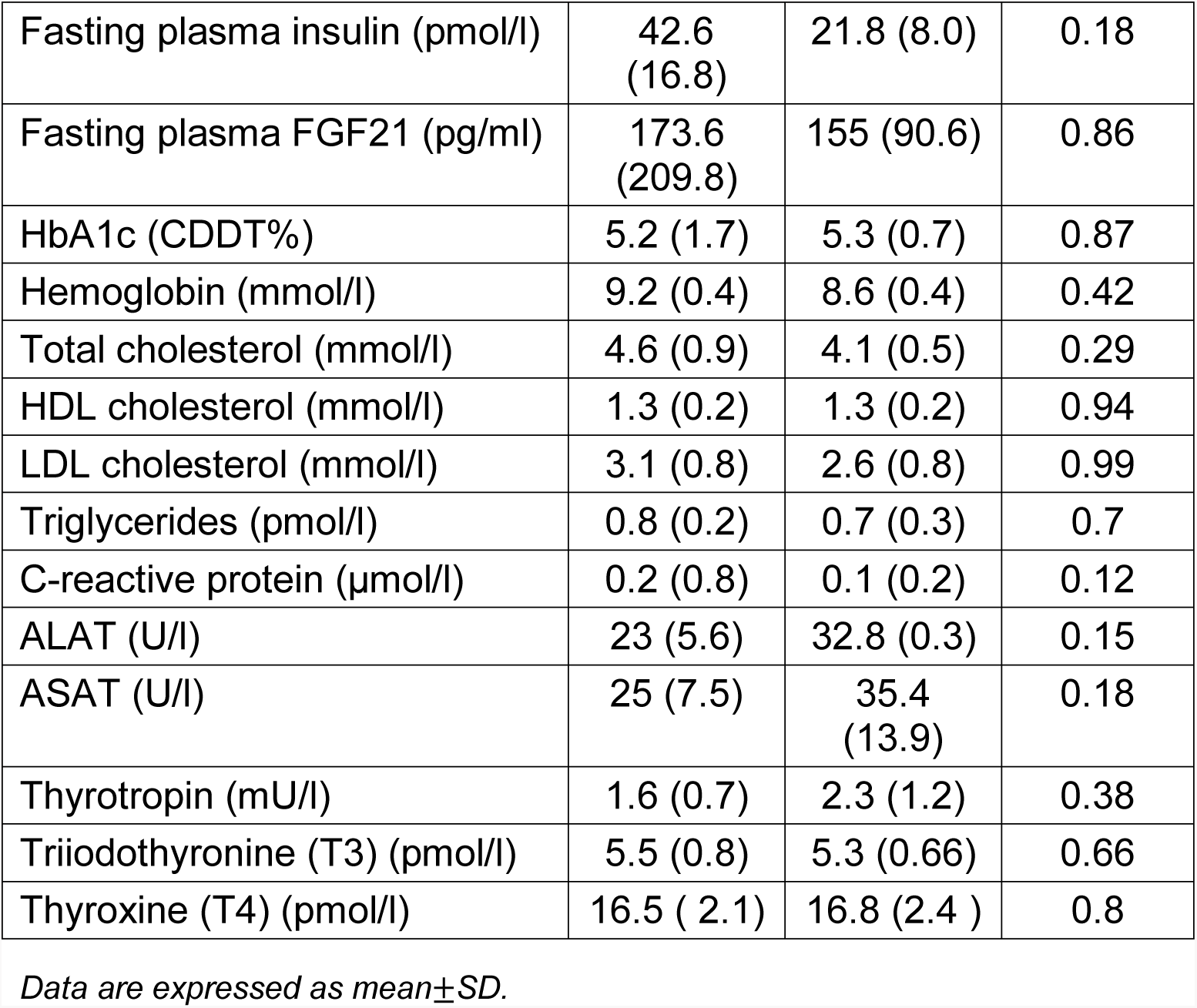
Subject characteristics.

|  | Alcohol<br>(n=5) | Water<br>(n=5) | *p* value |
| --- | --- | --- | --- |
| Age (years) | 24 (4.9) | 27 (1.5) | 0.26 |
| Body mass index (BMI) (kg/m) | 23.4 (1.6) | 24.4 (1.6) | 0.39 |
| Waist circumference (cm) | 84.2 (5.6) | 83.3 (6.6) | 0.84 |
| Fasting plasma glucose<br>(mmol/l) | 4.5 (0.2) | 4.5 (0.3) | 0.31 |
| Fasting plasma insulin (pmol/l) | 42.6<br>(16.8) | 21.8 (8.0) | 0.18 |
| Fasting plasma FGF21 (pg/ml) | 173.6<br>(209.8) | 155 (90.6) | 0.86 |
| HbA1c (CDDT%) | 5.2 (1.7) | 5.3 (0.7) | 0.87 |
| Hemoglobin (mmol/l) | 9.2 (0.4) | 8.6 (0.4) | 0.42 |
| Total cholesterol (mmol/l) | 4.6 (0.9) | 4.1 (0.5) | 0.29 |
| HDL cholesterol (mmol/l) | 1.3 (0.2) | 1.3 (0.2) | 0.94 |
| LDL cholesterol (mmol/l) | 3.1 (0.8) | 2.6 (0.8) | 0.99 |
| Triglycerides (pmol/l) | 0.8 (0.2) | 0.7 (0.3) | 0.7 |
| C-reactive protein (μmol/l) | 0.2 (0.8) | 0.1 (0.2) | 0.12 |
| ALAT (U/l) | 23 (5.6) | 32.8 (0.3) | 0.15 |
| ASAT (U/l) | 25 (7.5) | 35.4<br>(13.9) | 0.18 |
| Thyrotropin (mU/l) | 1.6 (0.7) | 2.3 (1.2) | 0.38 |
| Triiodothyronine (T3) (pmol/l) | 5.5 (0.8) | 5.3 (0.66) | 0.66 |
| Thyroxine (T4) (pmol/l) | 16.5 ( 2.1) | 16.8 (2.4 ) | 0.8 |

Data are expressed as mean±SD.

In baseline samples (t=0) from these subjects, we observed substantial variation (9-490 pg/ml) in FGF21 (1-181) between individuals, irrespective of group (Table 1). Accordingly, FGF21 results are expressed as fold-changes from baseline for each subject, to express treatment effects. As expected, ingestion of water did not lead to a change in levels of active FGF21 (1-181) at any time point, whereas ingestion of alcohol resulted in a 2.3-fold increase in circulating FGF21 (1-181) levels after 90 minutes, with a peak 3.4-fold over baseline at 120 minutes (Figure 1A).

### 3.2 Bolus treatment with recombinant human FGF21 reduced alcohol intake in mice

We subsequently investigated the potential physiological role of this transient increase in FGF21 after acute alcohol ingestion in mice. After a four-day baseline period with unrestricted, simultaneous access to water and a 4% ethanol solution, mice were injected with recombinant human FGF21 or vehicle once per day, and their *ad libitum* alcohol and water intake was compared to their baseline intake. This design was used to mitigate inter-individual differences caused by variable but stable preferences for a specific ratio of water to alcohol consumed by each mouse. In agreement with similar reports using infusion minipumps, FGF21-treated mice drank 21% less 4% alcohol solution relative to their own, untreated baseline (Figure 1B), and this decrease was statistically significant (p=0.015). By contrast, vehicle control-treated mice consumed the same amount of the 4% alcohol solution during treatment relative to baseline (2.2 ± 0.8 ml/day at baseline vs. 1.94 ± 0.7 ml/day with vehicle treatment, n=9, p=0.12). In addition, water intake was similar before and after treatment with vehicle (2.6 ± 0.8 ml/day pre vs. 2.4 ± 0.6 ml/day post, n=9, p=0.54), or FGF21 (2.2 ± 0.8 ml/day pre vs.2.5 ± 0.8 ml/day post, n=8, p=0.39).

**Figure 1.**
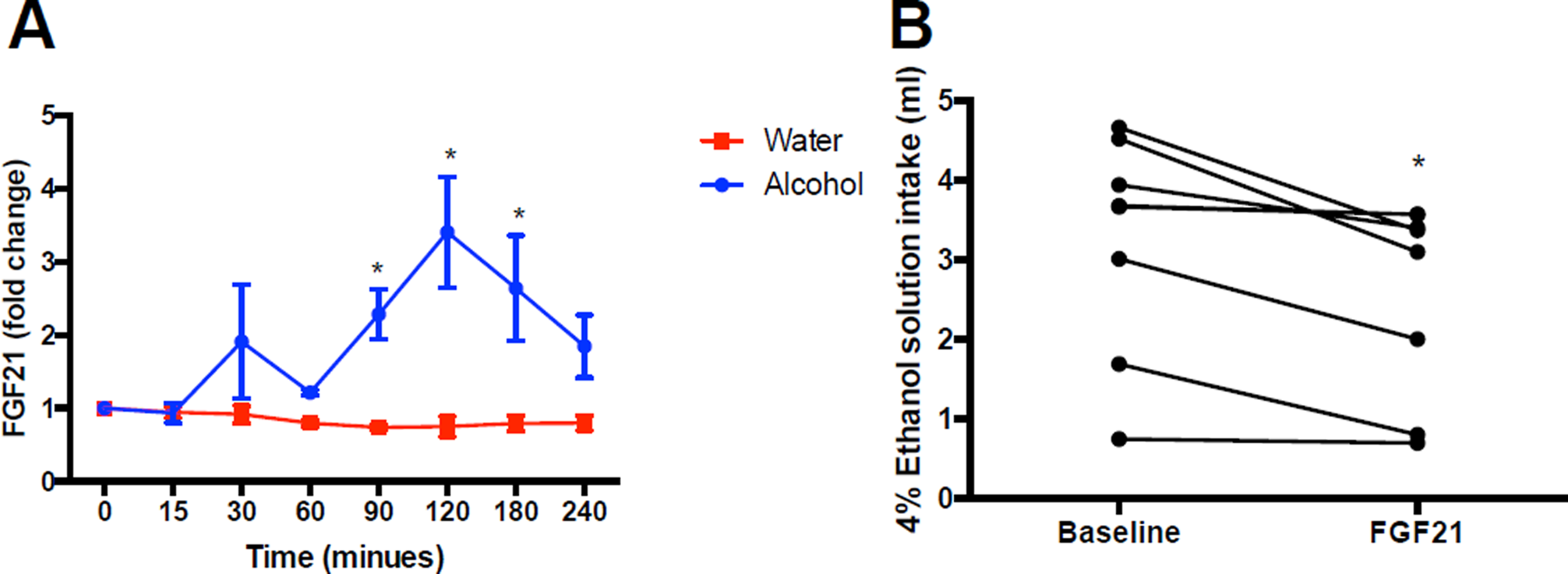
**A)** Relative fold change in active plasma FGF21 (1-181) in humans after drinking water or ethanol in water (≈2 standard US drinks/3 UK units), normalized to FGF21 levels at t=0 for each subject (n=5/group, ∗ denotes statistical significance where Benjamani-Hochberg Q<0.05. **B)** Effect of intraperitoneal FGF21 administration (1 mg/kg) on average daily intake of a 4% ethanol solution in mice over four days of baseline and three days of FGF21 treatment (n=8-9/group), ∗ Denotes statistical significance, p<0.015 by paired, two-tailed Student’s t-test.

### 3.3 Oktoberfest binge drinking increased total plasma FGF21 but produced relatively minor changes in hepatic and metabolic markers

At the festival, the three subjects consumed an average of 22.6 beers/person/day (4.4 g/kg/day pure ethanol or 7.46 Lbeer/person/day) (range=16-28 beers/day) for three days. Remarkably, the three-day period of concerted drinking elicited relatively minor changes in blood chemistry, and no changes in liver stiffness and triglyceride content assessed by fibroscan. As expected, plasma levels of ASAT, a marker of hepatic injury, increased by an average of 25% relative to baseline measurement, but remained within the normal clinical range. Another circulating marker of hepatocyte lysis, ALAT, tended to increase to a similar extent, but also remained within normal clinical limits (Supplementary Table 4). With respect to metabolic parameters, fasting plasma glucose was higher in all subjects after the return from the Oktoberfest compared to baseline, but was not accompanied by changes in insulin, C-peptide, HOMA-IR or HOMA-2-IR. However, plasma triglyceride increased by an average of 161% following Oktoberfest exceeding the normal clinical range in all three subjects (Supplementary Table 4). Similarly, plasma FGF21 (1-181) was elevated by an average of 110% upon return (Figure 2), and total plasma FGF21, including cleavage products, was increased 90% (Supplementary Table 4).

**Figure 2.**
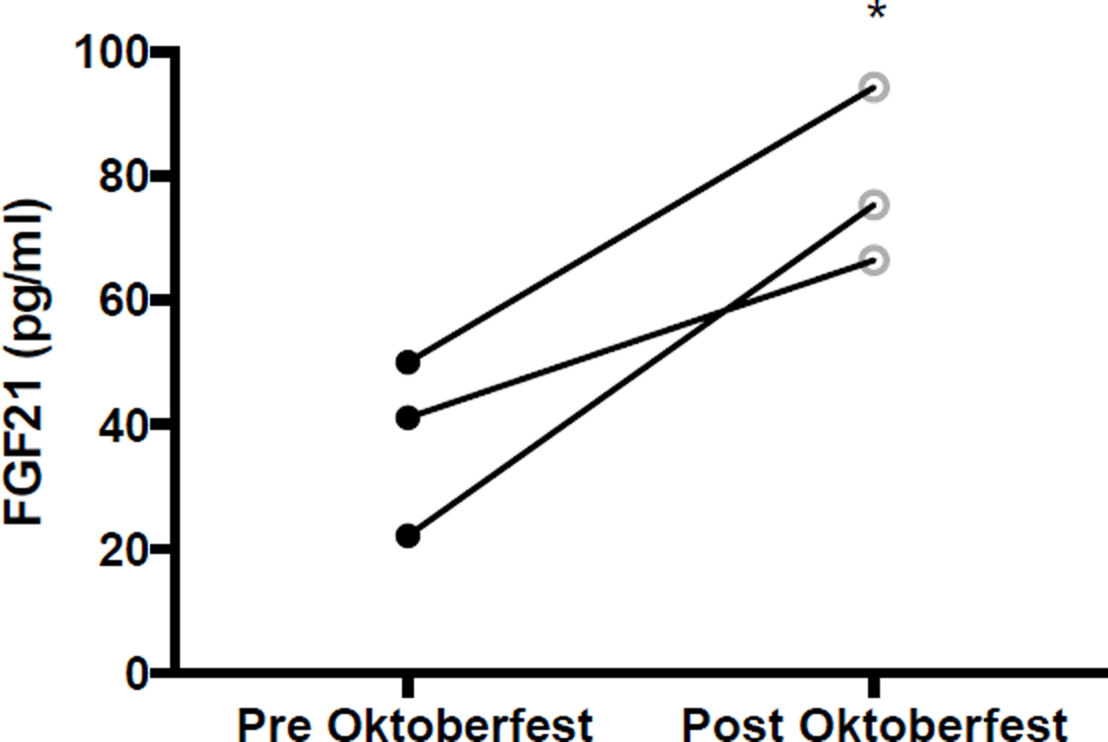
Active plasma FGF21 (1-181) levels before and after consuming an average of 22.6 beers/day for three days at Oktoberfest 2017. ∗ Denotes statistical significance, p<0.05 by paired, two-tailed Student’s t-test.

### 3.4 Basal levels of active FGF21 (1-181) in human plasma were not associated with alcohol-related behavior or problems in non-alcoholic humans

Together, our results in humans and mice suggest that FGF21 (1-181) may increase after alcohol drinking to discourage further alcohol consumption. Therefore, we investigated whether FGF21 (1-181) levels are correlated with alcohol drinking patterns or problems in humans. Subjects from the alcohol challenge study described above (n=10), as well as participants from a previous study by our group that investigated FGF21 and sweet preference (n=39), completed two alcohol use questionnaires: AUDIT and Alcohol E (see Materials and Methods for details). Because the two cohorts were similar in demographic and anthropometric characteristics (Supplementary Table 2), their responses were pooled for analysis. For each survey question and the total questionnaire score, we calculated a Spearman correlation coefficient (r) with fasted levels of active plasma FGF21 (1-181). However, we did not observe a significant relationship between FGF21 levels at baseline and any alcohol-related behaviors, emotional responses, or problems in either AUDIT or Alcohol E in this small cohort (p>0.05 for all correlations) (Supplementary Table 3).

## 4. DISCUSSION

Here we show, in humans, that FGF21 (1-181) levels in circulation are increased more than 3-fold by acute alcohol intake and that sub-chronic alcohol binge drinking in a natural setting (Oktoberfest, Munich) persistently increases FGF21 levels. Further, we show that once-daily bolus injection of recombinant human FGF21, in mice, is sufficient to reduce *ad libitum* alcohol intake by 21%.

Our findings support the negative-feedback model of Schumann et al., who associated variants in *KLB* with alcohol intake in humans, and further demonstrated that continuous FGF21 infusion by minipump reduces alcohol intake through β-klotho in the central nervous system [4]. Here we provide evidence consistent with their prediction that the human liver secretes FGF21 in response to alcohol intake. Our work is also consistent with a recent report by Desai and coworkers that alcohol acutely increases total circulating FGF21 in non-fasting humans [22]. While we observed a smaller increase in circulating FGF21 after alcohol ingestion when compared to their 0.9 g/kg dose (3.4-fold vs 40-fold), the explanation for this may be a non-linear relationship between FGF21 secretion and increasing doses of alcohol. The dose we used was slightly lower than the lowest 0.4 g/kg dose in their study, which evoked less FGF21 secretion than the 0.9 g/kg dose (10-fold vs. 40-fold). Thus, alcohol bingeing may elicit exaggerated FGF21 secretion relative to moderate drinking, which could be relevant to the pathogenesis of alcohol abuse if FGF21 has reinforcing, in addition to acute anti-consumptive, effects in the central nervous system. In addition, because alcohol appears to be the strongest physiological inducer of FGF21 identified to date in humans, it will be of interest to uncover the mechanism responsible for alcohol-mediated FGF21 secretion, as it may offer new targets for stimulating FGF21 production to achieve therapeutic ends.

Another potential contributor to the difference in magnitude and temporal pattern of FGF21 secretion between our study and that of Desai *et al.* is the fact that their subjects had access to food, which has complex effects on FGF21 production in humans, whereas our subjects were fasted [19, 23, 24]. A further salient difference between our acute studies was the ELISA assay kit used;

Desai *et al.* measured total FGF21 whereas we measured only active, full-length FGF21 in our acute study. We focused on active FGF21 in light of recent reports that FGF21 signaling is terminated by FAP-mediated cleavage at the C-terminus, and that C-terminally intact FGF21 is highly correlated with total FGF21 when total FGF21 is measured with a capture antibody targeted to avoid regions subject to proteolysis [25]. Therefore, one reason Desai et al. may have identified a later peak in FGF21 than we did is due to the accumulation of FGF21 fragments. However, future studies will be needed to distinguish this possibility from an augmented and delayed peak in FGF21 caused by the admixture of food and alcohol.

FGF21 overexpression or continuous infusion by minipump decreases alcohol intake in mice [4, 13]. However, it was not clear whether bolus injection of FGF21, whose t_1/2_ is approximately 1-2 hours [26], would have long-lasting effects on alcohol intake. Because FGF21 (1-181) peaks and declines rapidly after alcohol drinking, and because we previously observed a durable effect of bolus FGF21 administration on sweet appetite [27], and suspect that this ability to reduce sweet and alcohol intake may be mediated by a partially overlapping population of β-klotho positive neurons, we hypothesized that bolus injection of FGF21 (1-181) would also reduce alcohol intake over a 24 hour period. Consistent with this, human FGF21 administered once every 24 hours was sufficient to reduce overall alcohol intake by 21% in mice. This suggests that transient increases in FGF21 may have persistent effects on drinking behavior.

Our results are reminiscent of prior reports that oral boluses of fructose [20], sucrose [15], and glucose [21] acutely increase circulating FGF21 levels in humans and rodents. In these cases, the change in FGF21 secretion is smaller and may be a physiological, negative-feedback mechanism to inhibit sweet intake; *Fgf21* knockout mice consume twice as much sucrose as wild type [27], FGF21 (or FGF21 analogue) injection reduces sweet intake in mice and nonhuman primates [13, 27], and the rs838133 A-allele in the *Fgf21* gene is associated with increased sweet snacking in humans [15]. These studies provide evidence that FGF21 reduces sweet intake in a physiological context. Given our current data, we posit that there may be an overlapping circuit regulating alcohol intake that also relies on FGF21. Indeed, we previously reported that the FGF21 rs838133 A-allele is associated with increased alcohol consumption in the Danish Inter99 cohort (n=6,514) [15]. However, in contrast to sugars, *Fgf21* knockout mice do not consume more alcohol than wild type, leaving the physiological role of FGF21 in regulating alcohol intake in doubt [22]. One possible explanation for this is compensatory signaling via FGF15/19, though this remains speculative.

Intriguingly, although pharmacological FGF21 treatment reduces body weight and food intake in primates [19], the sweet-increasing *Fgf21* rs838133 A-allele is linked to slightly lower BMI in humans. This is at odds with the concept that FGF21 regulates body weight by limiting sweet and alcohol intake [15]. An alternative hypothesis is that a negative-feedback axis between the liver and brain evolved to limit overindulgence in fructose and alcohol from fermented fruits and nectars, as this can be acutely hepatotoxic. For example, the pentailed treeshrew *(Ptilocerus lowii),* a mammal similar to an extinct primate ancestor of modern humans, consumes large amounts of fermented nectar from the bertam palm *(Eugeissona tristis),* which has alcohol content similar to beer. Thus, human ancestors may have experienced pressure to balance the value of alcohol as an energy source with its deleterious effects on liver when consumed in excess [28]. The ability of FGF21 to stimulate hepatic fatty acid oxidation may be one example of such an adaptation [8]. Another possibility is that the induction of FGF21 by dietary sugars and alcohol may promote balanced food selection, in particular protein intake. Indeed, FGF21 is strongly upregulated by protein restriction [29, 30], suggesting that limiting the drive to consume foods that are calorie-rich but protein poor may be adaptive after a certain threshold of intake.

Our finding that three days of heavy drinking at Oktoberfest increased fasting plasma FGF21 (1-181) levels by 110% after 32 hours of alcohol abstinence raises the possibility that FGF21 may regulate alcohol intake in both sub-chronic and acute contexts. However, the increase in FGF21 after Oktoberfest may also be a response to increased carbohydrate intake and hepatic lipid content derived from excess food consumption, which has previously been demonstrated in humans, and is also consistent with the large increases in circulating triglyceride observed in these subjects [23]. Therefore, further studies will be required to establish a direct link between increased sub-chronic levels of alcohol intake and changes in circulating FGF21 in humans. In addition, it is notable that FGF21 is increased dramatically relative to canonical liver enzymes (e.g. AST and ALT) by binge drinking, suggesting that it may be a promising biomarker for recent alcohol abuse. Finally, while one might conclude that the FGF21 induction caused by short-term alcohol intake reflects transient hepatic stress, recalling that FGF21 secretion is also elevated in mitochondrial myopathies [31], in fact the hepatotoxin acetaminophen decreases circulating FGF21 levels [32], more consistent with the idea that FGF21 secretion may be selectively regulated by alcohol ingestion to fulfill a signaling function (e.g. stimulate fatty acid oxidation or promote alcohol satiety).

Given the genetic link between FGF21 and alcohol intake in humans, it is notable that we did not find a correlation between fasting FGF21 (1-181) levels in plasma and alcohol-related behavior, emotional responses, and problems as measured by the AUDIT and ALCOHOL-E questionnaires. Potential explanations for this are the fact that our sample size was small, and did not include subjects with abnormal drinking behavior. Otherwise, the association between *KLB* variants and alcohol intake may be mediated by FGF19, which comes from the gastrointestinal tract and can also signal via the FGFR1/β-klotho receptor in the brain, in addition to the FGFR4/β-klotho complex in the liver to regulate bile acid production [33]. Future studies will be required to distinguish between these alternatives.

In conclusion, the present work supports the hypothesis that circulating FGF21 may be a liver-derived inhibitor of alcohol intake in humans, akin to gut peptides that reduce food intake after meals. However, further work is required to establish this via clinical trials and to understand whether hepatic hormones play a more general role than anticipated in regulating ingestive behavior, given the centrality of the liver as a metabolic integrator.

## ACKNOWLEDGEMENTS

The authors gratefully acknowledge the participants in this study as well as colleagues, in particular Cecila Ratner Christoffer Clemmensen, for helpful comments and discussion. We also thank Birgitte Andersen (Novo Nordisk) for providing recombinant FGF21. This work was supported by the Novo Nordisk Foundation Center for Basic Metabolic Research. The Novo Nordisk Foundation Center for Basic Metabolic Research is an independent Research Center at the University of Copenhagen partially funded by an unrestricted donation from the Novo Nordisk Foundation (http://metabol.ku.dk/). The Centre for Physical Activity Research is supported by a grant from TrygFonden. The Centre of Inflammation and Metabolism/ Centre for Physical Activity Research is a member of DD2, the Danish Center for Strategic Research in Type 2 Diabetes (the Danish Council for Strategic Research; Grants 09 - 067009 and 09 - 075724).

